# Axonal mTOR–dependent Rab5 translation drives axonal transport and BDNF signaling to the nucleus

**DOI:** 10.1101/2025.10.14.682461

**Authors:** Reynaldo Tiburcio-Felix, Constanza Tapia-Peralta, Pablo Ahumada-Montalva, Gloria Arriagada, Eran Perlson, Francisca C. Bronfman

**Author notes:** These authors contributed equally to this work (Reynaldo Tiburcio-Felix, Constanza Tapia-Peralta).

## Abstract

Brain-derived neurotrophic factor (BDNF) promotes neuronal plasticity through retrograde signaling from axon terminals to the nucleus, activating CREB-dependent transcription. While this relies on signaling endosomes, the molecular mechanisms enabling their axonal transport remain poorly understood. Using compartmentalized cultures of mouse cortical neurons, we show that axonal BDNF activates mTOR signaling and stimulates local protein synthesis. Translation inhibitors blocked both BDNF-enhanced retrograde transport and nuclear CREB phosphorylation, indicating that axonal protein synthesis is required for long-distance signaling. Among locally synthesized proteins, we identified Rab5, a master regulator of endosomal trafficking. BDNF increased axonal Rab5 levels through TrkB- and mTOR-dependent mechanisms, confirmed by puromycin-PLA for Rab5 and soma-free axon preparations. Axon-specific Rab5 knockdown abolished BDNF-induced retrograde transport and CREB activation, demonstrating that local axonal translation of Rab5 mRNA is essential for neurotrophin signaling propagation. Remarkably, even basal retrograde transport depended on ongoing axonal Rab5 synthesis, revealing a constitutive role for local translation in maintaining axonal trafficking capacity. These findings establish that local axonal translation of trafficking regulators is a prerequisite for axon-to-nucleus neurotrophin signaling, positioning on-demand protein synthesis as a central node in long-distance neuronal communication.

## INTRODUCTION

Neuronal plasticity, the ability of neurons to adapt and reorganize, is fundamental to learning and memory and is driven by structural changes in synapses, including dendritic and axonal remodeling, regulated by neuronal activity and neurotrophic factors (NTFs). Among NTFs, brain-derived neurotrophic factor (BDNF) is the most extensively studied in the central nervous system (CNS). Secreted in an activity-dependent manner, BDNF activates its receptor, tropomyosin receptor kinase B (TrkB), inducing local cytoskeletal changes and long-distance signaling that influence nuclear gene expression, thereby supporting synaptic remodeling and circuit maintenance^1–3^.

Recent studies, including our own, have shown that BDNF signaling in cortical axons generates specialized signaling endosomes containing BDNF and activated TrkB^4–6^. These endosomes facilitate axon-to-nucleus communication, promoting CREB-dependent transcription of plasticity-related genes such as Arc and enhancing dendritic branching. This mechanism highlights the critical role of endosomal trafficking in maintaining long-distance neurotrophin signaling.

Rab GTPases are key regulators of membrane trafficking, orchestrating the dynamics of signaling receptors. Acting as molecular switches cycling between GDP- and GTP-bound states, Rab proteins recruit effectors to specific membrane compartments. Rab5 mediates the first step of receptor sorting and vesicle fusion at early endosomes, Rab11 facilitates receptor recycling, and Rab7 governs the transition to late endosomes^7–9^. Our previous work showed that BDNF enhances Rab5 and Rab11 activity in hippocampal neurons and that reduced Rab function impairs BDNF-induced dendritic arborization^10–12^, linking signal transduction to endosomal trafficking.

Rab5 also regulates long-distance neurotrophin signaling in axons. In motoneurons, Rab5 facilitates the early endosomal sorting of TrkB^13^, and in sympathetic neurons, its activity is linked to BDNF-signaling endosomes that mediate retrograde cell death signals^14^. In cortical neurons, Rab5-mediated retrograde transport is regulated by Rab5 effectors^15,16^, placing Rab5 at the intersection of endosomal sorting and motor-driven retrograde transport. Together, these findings establish Rab5 as a conserved regulator of retrograde neurotrophin signaling across neuronal subtypes.

BDNF binding to TrkB activates multiple downstream cascades, including ERK1/2, PI3K-AKT, and PLCγ-Ca2+, leading to CREB activation and dendritic arborization^1,17^. Among these, the mechanistic target of rapamycin (mTOR) pathway is a central regulator of activity-dependent protein synthesis. mTOR promotes translation initiation through phosphorylation of 4E-binding proteins (4EBPs) and ribosomal S6 kinases (S6Ks)^18^, and is well-established as a key mediator of BDNF-induced translation in dendritic spines^19^.

Local protein synthesis in axons and dendrites enables rapid, spatially precise responses to extracellular stimuli and is essential for neuronal development and function. Disruptions in local proteostasis are linked to neurodegenerative and neurodevelopmental diseases^20–22^. Importantly, local translation in axons has been shown to regulate the availability of trafficking components, suggesting that on-demand protein synthesis could directly influence axonal transport capacity^23–25^. Notably, small GTPases, including TC10, RhoA, and Cdc42, undergo local translation in growth cones and axons^26–28^, and our previous work showed that BDNF enhances Rab5 protein levels and activity in hippocampal neurons in an mTOR-dependent manner^10^, raising the possibility that Rab5 may itself be a target of axonal local translation downstream of BDNF-mTOR signaling. However, whether BDNF activates mTOR-dependent translation specifically within axons and whether this contributes to retrograde transport and nuclear signaling remain unknown.

Here, using compartmentalized cultures of mouse cortical neurons, we demonstrate that axonal BDNF stimulation activates mTOR signaling and drives local protein synthesis. We identify Rab5 as a key locally translated protein whose synthesis is required for BDNF-enhanced retrograde transport and nuclear CREB activation, revealing a previously unrecognized link between axonal proteostasis and long-distance neurotrophin signaling and establishing local Rab5 synthesis as a new regulatory node linking axonal mTOR activation to axonal transport and propagation of BDNF signaling to the nucleus.

## RESULTS

### BDNF in axons of cortical neurons increases local protein synthesis in a mTOR-dependent manner

Multiple lines of evidence indicate that BDNF activates mTOR in dendrites and spines^2,29^. However, the functional role of axonal local protein translation mediated by BDNF-TrkB-mTOR in CNS neurons has not been addressed. To investigate whether BDNF increases mTOR activity in axons, we employed compartmentalized cortical neuron cultures using microfluidic devices, a model that we and others have extensively validated^6,14,30,31^. Axons were divided into three regions: microgrooves (MG), proximal axons (PA), and distal axons (DA) (Figure 1A, B). To restrict BDNF signaling to the axonal compartment (AC), endogenous BDNF in the cell body compartment (CB) was neutralized with TrkB-fc^32^; also, the media flux was maintained towards the axonal compartment, and compartmentalization was confirmed using fluorescent cholera toxin B (f-Ctb), which undergoes retrograde transport to label connected cell bodies (Figure 1A, B).

**Figure 1.**
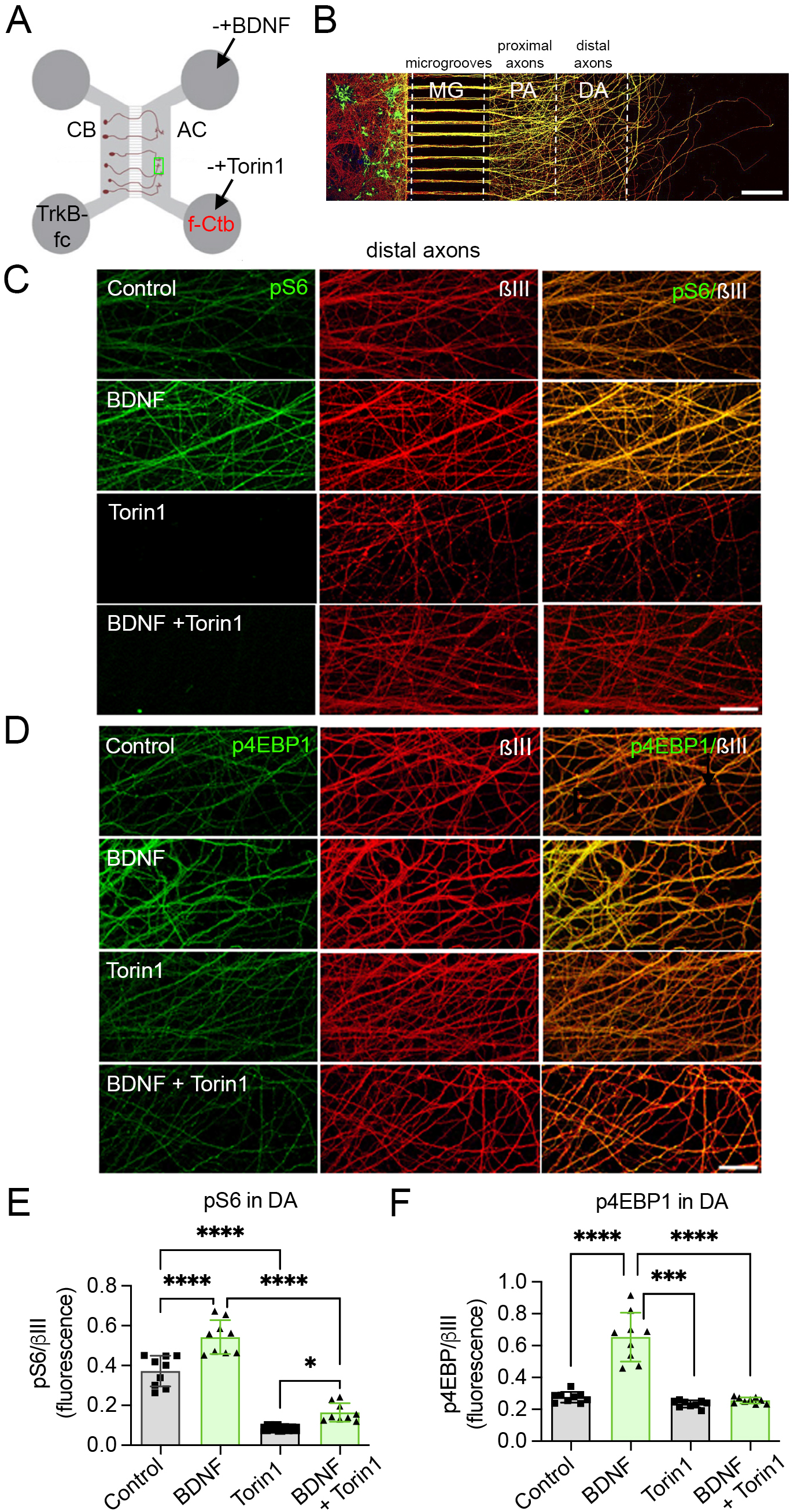
BDNF induces mTOR-dependent phosphorylation of ribosomal protein S6 (pS6) and eukaryotic translation initiation factor 4E-binding protein 1 (4EBP1) in axons of compartmentalized cortical neurons. **A.** Experimental design and diagram of compartmentalized cultures of mouse cortical neurons in microfluidic chambers, used to study local BDNF-induced signaling. The cell body (CB) and axonal compartments (AC) are labeled. On DIV6, 1 µg/mL of fluorescent cholera toxin subunit B (f-Ctb) was added to the AC. The next day, TrkB-fc (100 ng/mL) was added only to the cell bodies for 60 minutes. Subsequently, BDNF (50 ng/mL) was applied to the axons for 180 minutes. **B**. Representative image of compartmentalized cultures on DIV7. βIII-tubulin (red) and f-Ctb (green) were detected by confocal microscopy. The cell body compartment (CB), microgrooves (MG), proximal axons (PA), and distal axons (DA) are distinguished. Scale bar, 200 µm. **C and D**. Cortical neurons in compartmentalized cultures were treated with or without BDNF (50 ng/mL) and with or without the mTOR inhibitor Torin1 (0.25 µM). Fixed cultures were labeled for pS6 (green) in **C** or p4EBP1 (green) in **D**, with βIII-tubulin (βIII) in red. **E and F**. Quantification of immunofluorescence for pS6 in distal axons (DA) in **E** and p4EBP1 in **F**. Conditions include non-treated axons (control), axons treated with BDNF (BDNF), axons treated with Torin1 (Torin1), and axons treated with BDNF in the presence of Torin1 (BDNF + Torin1). Data represent the mean ± SD from n=9-10 chambers across three independent experiments. Statistical analysis was performed using one-way ANOVA and Tukeýs multiple comparison test (*p<0,05; ***p<0,001; ****p<0,001).

Axonal BDNF treatment significantly increased phosphorylation of the mTOR targets pS6 and p4EBP1 along the axon (Figure 1C-F), an effect abolished by the mTOR inhibitor Torin1^33^ (Figure 1C-F), confirming that BDNF specifically activates the mTOR pathway in axons.

To determine whether mTOR activation led to functional protein synthesis, we treated axons with BDNF in the presence of O-propargyl-puromycin (OPP), which incorporates into nascent peptides^34^. Using Click-it chemistry to visualize newly synthesized proteins, we observed a 2-fold increase in protein synthesis following BDNF treatment. This increase was prevented by both Torin1 and cycloheximide (CHX), a protein synthesis inhibitor^19,35^. Thus, BDNF triggers mTOR-dependent protein synthesis in axons (Figure 2).

**Figure 2.**
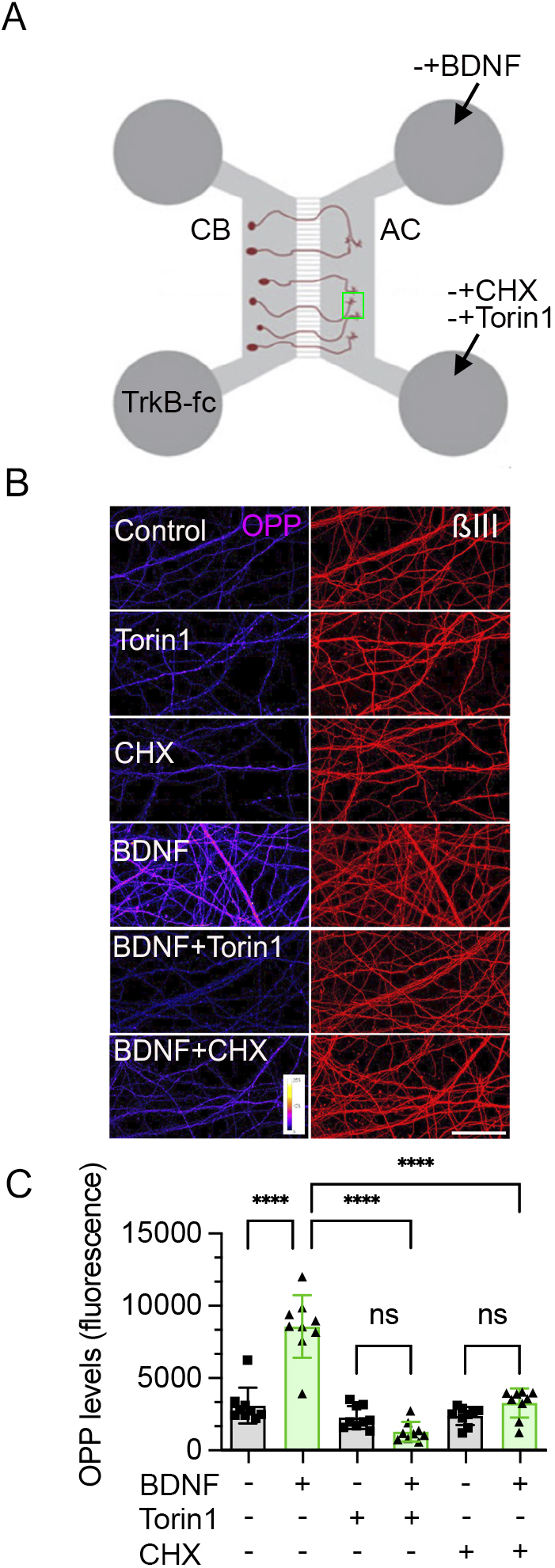
BDNF induces axonal protein synthesis in a mTOR and translation-dependent manner. **A.** The AC was treated with BDNF or vehicle (control) in the absence or presence of Torin1 (0,25 µM) and cycloheximide (CHX, 25 µM) for 180 minutes in the presence of O-propargyl-puromycin (OPP). **B.** OPP was visualized using click-it chemistry with a picolyl azide conjugate Alexa Fluor^TM^ 488 dye and visualized using a magenta scale, as shown in the insets (last left panel). Axons were co-stained with an antibody against βIII-tubulin (βIII) (right panels), shown in red. **C.** Quantification of OPP fluorescence intensity (arbitrary units). Values represent mean ± SD of n=9-10 chambers from three independent experiments. Statistical analysis was performed using one-way ANOVA and Tukeýs multiple comparison test (****p < 0.0001, ns, not significant).

### Local protein synthesis induced by BDNF in cortical neurons is required for axonal transport and CREB phosphorylation

We previously reported that BDNF enhances axonal retrograde transport of f-Ctb in cortical neurons^30^. Consistent with this, co-application of f-Ctb and BDNF for 3 hours doubled f-Ctb transport to somas (Figure 3A-C). Significantly, this enhanced transport was abolished by anisomycin, a protein synthesis inhibitor^36^ (Figure 3B and 3C), indicating that BDNF-induced retrograde transport requires newly synthesized axonal proteins. f-Ctb co-traffics with BDNF/TrkB signaling endosomes^6,30,37^, and retrograde transport of these endosomes enhances nuclear CREB phosphorylation, leading to plasticity gene expression and dendritic arborization^6^. To test whether axonal protein synthesis is required for this nuclear signaling, we compared pCREB levels in neurons treated with BDNF alone or with BDNF plus CHX in the AC (Figure 3D). CHX significantly reduced nuclear pCREB levels (Figure 3D, E), demonstrating that local axonal translation is essential for BDNF-induced nuclear signaling.

**Figure 3.**
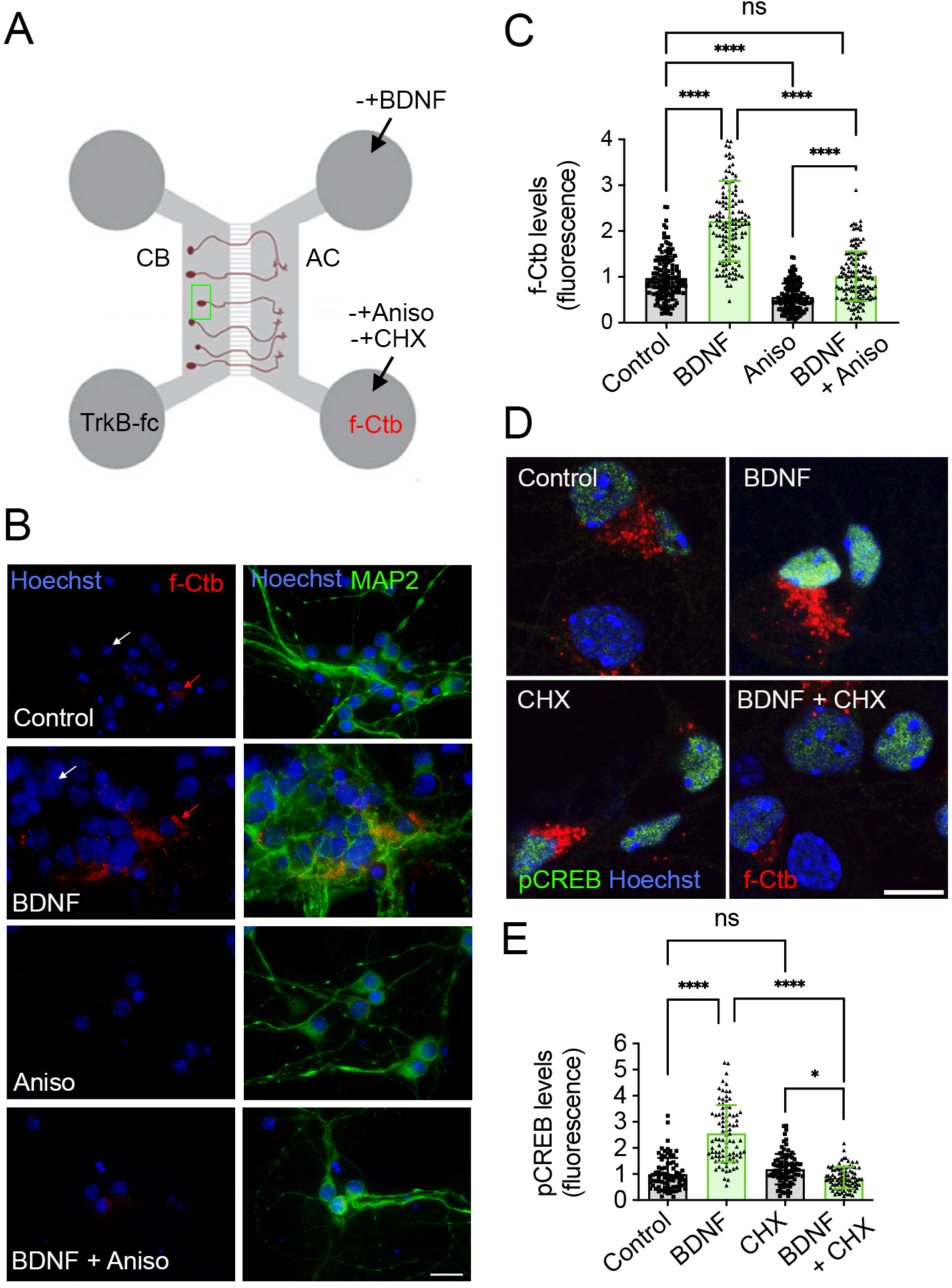
Axonal BDNF increases the retrograde transport of f-Ctb and phosphorylated levels of CREB, a process regulated by axonal local translation. **A.** Cortical neurons were cultured in microfluidic chambers up to day 7 (DIV7). Then, TrkB-fc (100 ng/mL) was added for 60 minutes of treatment in the CB compartment; the AC was treated or not with 25 µM anisomycin (Aniso), in the case of experiments evaluating f-Ctb transport (B and C) or 25 µM cycloheximide (CHX), in the case of experiments evaluating pCREB levels (D and E) for 1 hour before treating axons with or without BDNF (50 ng/mL) for 180 minutes in the presence of f-Ctb (1 µg/mL). **B.** Nuclei were stained with Hoechst (blue), and MAP2 immunostaining as a general neuronal marker was visualized in green. The right panel shows the overlap of the f-Ctb in red and Hoechst channels, and the left panel shows the overlap of the three labels. **C.** Quantification of the f-Ctb-associated fluorescence intensity in cell bodies relative to the control. The white arrows indicate neuronal somas without the f-Ctb label, and the red arrows indicate somas of a neuron with the f-Ctb mark. Only cells with a f-Ctb were considered in the quantification. Values represent mean ± SD of n=45-50 neurons from three independent experiments (3 different chambers per experiment). Statistical analysis was performed using one-way ANOVA and Tukeýs multiple comparison test (****p < 0.0001, ns, not significant). Scale bar, 20 μm. **D.** Nuclei were stained with Hoechst (blue) and immunostained for pCREB (Ser133), shown in green, in cells labeled with f-Ctb (red). Scale bar 10 µm. **E.** Quantification of nuclear pCREB fluorescence intensity. Values represent mean ± SD of 90 cells from three independent experiments. Statistical analysis was performed using one-way ANOVA and Tukeýs multiple comparison test (*p<0,05; ****p < 0.0001, ns, not significant).

### Axonal BDNF increases the local translation of Rab5 in a TrkB and mTOR-dependent manner

Our previous work showed that BDNF enhances Rab5 activity and protein levels in hippocampal neurons in an mTOR-dependent manner^10^. Given that mTOR-dependent local translation is required for BDNF-enhanced retrograde transport (Figures 2 and 3), we next asked whether Rab5 undergoes local translation in cortical axons in response to BDNF. Immunofluorescence analysis revealed that BDNF treatment doubled Rab5 levels in distal axons (Figure 4A-C). Using neurons from TrkBF616A mice, which express a kinase-domain mutation that allows pharmacological inhibition with 1NM-PP1^38^, we found that Rab5 upregulation was blocked by both 1NM-PP1 and Torin1 (Figure 4A-C), confirming dependence on TrkB and mTOR activity. CHX treatment also prevented Rab5 upregulation (Figure 4D and 4E), suggesting that the BDNF-induced increase in Rab5 results from de novo protein synthesis rather than from transport from cell bodies.

**Figure 4.**
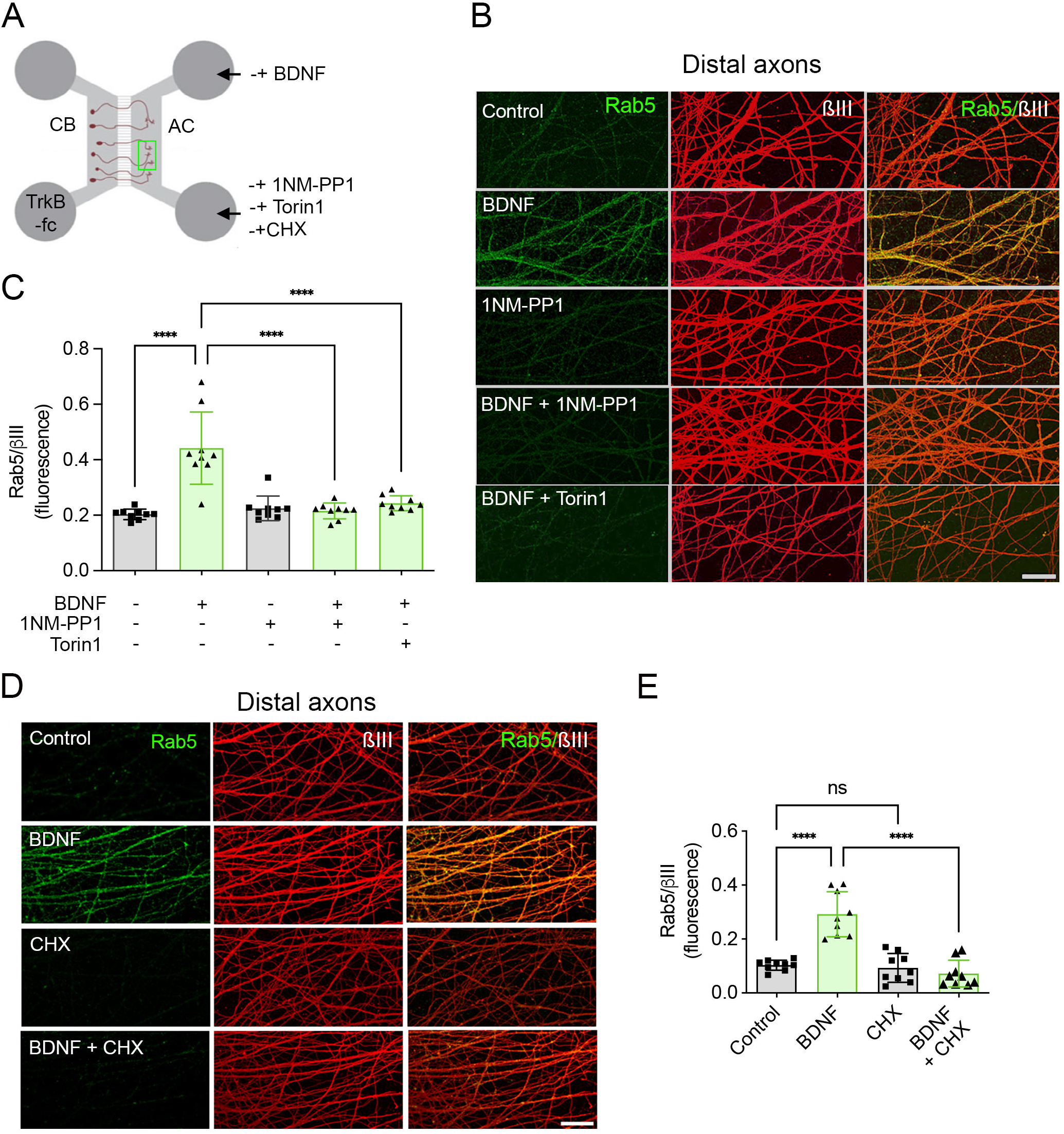
BDNF increases Rab5 levels in distal axons, a process that depends on axonal TrkB activity, mTOR, and protein translation. **A.** Cortical neurons were cultured in microfluidic chambers for 7 days. Then, TrkB-fc (100 ng/mL) was added to the CB compartment for 60 minutes of treatment. After, the AC was not treated (control) or treated in the presence of 1NM-PP1 (1 µM), CHX (25 µM), or BDNF (BDNF) alone or in the presence of 1NM-PP1, Torin 1 (0.25 µM), or CHX. **B.** Rab5 immunostaining in control axons or axons stimulated with BDNF or in the presence of TrkB and mTOR inhibitors (1NM-PP1 and Torin1, respectively). Neurons were labelled for axonal Rab5, shown in green, and βIII-tubulin (βIII), shown in red. Pictures were obtained by confocal microscopy in distal axons. **C.** Quantification of Rab5 immunofluorescence intensity in distal axons standardized by βIII-tubulin (βIII) immunostaining. **D.** Rab5 immunostaining in control axons or axons stimulated with BDNF in the absence of CHX for 180 minutes. Pictures were obtained by confocal microscopy in distal axons. Scale bar, 50µm**. E.** Quantification of Rab5 immunofluorescence intensity (green) in distal axons standardized by βIII-tubulin (βIII) immunostaining. Values represent the mean ± SD of n=9-10 chambers from three independent experiments. Statistical analysis was performed using one-way ANOVA and Tukeýs multiple comparison test (****p<0,0001; ns, not significant).

To confirm the axonal localization of Rab5 mRNA, we isolated RNA from the axonal and cell body compartments using radial chambers^39^ and performed RT-qPCR analysis. The quality of our axonal preparations was validated using an established marker: β-actin mRNA, known to be enriched in axons^40,41^, showed the expected axonal enrichment, whereas polymerase B mRNA, a nuclear-localized transcript, was detected only in cell body compartments and was absent from axonal preparations. Under these validated conditions, Rab5 mRNA was detected in both axonal and cell body compartments (Figure 5A-C), confirming its presence in axons. Western blotting of protein lysates confirmed that axonal lysates were devoid of nuclear markers and enriched in axonal markers (Figure 5E). Importantly, western blot analysis of axonal proteins indicated that BDNF treatment increased Rab5 protein levels, an effect blocked by CHX (Figure 5D-G).

**Figure 5.**
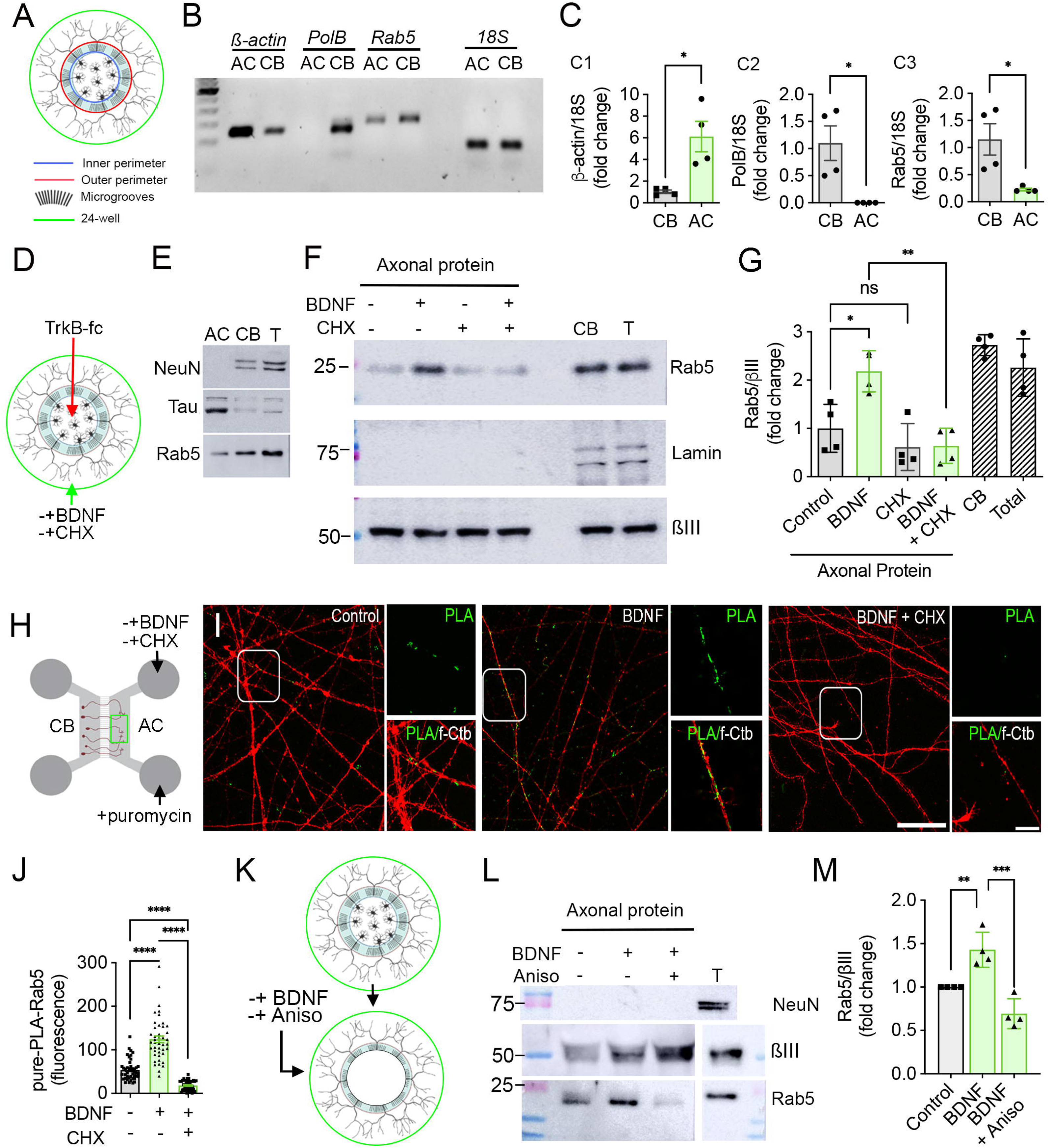
BDNF increases Rab5 mRNA translation in axons. **A**. Scheme of a radial chamber used to obtain axonal (AC) and cell body (CB) lysates of cortical neurons. **B**. Cortical neurons were cultured in radial compartmentalized cultures up to day 7. Then, lysates obtained from the axonal compartment (AC) were analyzed by RT-qPCR using primers for Rab5 (targeting all Rab5 isoforms), β-actin, polymerase B (PolB), and 18S RNA. The bands correspond to RT-qPCR products. **C.** RT-qPCR-based mRNA quantification of *ß-actin* in C1, *PolB* in C2, and *Rab5* in C3, relative to cell body levels of each gene. *18S* RNA was used as a housekeeping gene. Values represent the mean ±SEM of four independent experiments. Statistical analysis was performed by Unpaired T-Student (*p<0,05). **D.** Cortical neurons were cultured in radial compartmentalized cultures up to day 7. Then, TrkB-fc (100 ng/mL) was added to the CB compartment for 60 minutes of treatment. After, the AC was not treated (control) or treated in the presence of CHX, BDNF or BDNF including CHX (BDNF +CHX). **E.** RIPA protein lysates from AC or CB were characterized by the presence of a nuclear marker (NeuN), an axonal marker (Tau), and Rab5. T, indicates total lysate, including cell bodies and axons. **F.** Cortical neurons were cultured in radial compartmentalized cultures for 7-8 days. Then, neurons were treated as described in D, and lysates were obtained from the AC (axonal proteins), CB, and total proteins (T). The levels of Rab5, Lamin A/C (nuclear marker), and βIII-tubulin (βIII) were assessed by western blotting. CB or T lysates were untreated. **G.** Quantification of axonal Rab5 normalized by βIII protein levels. Values represent the mean ±SD, n=4. one-way ANOVA and Tukeýs multiple comparison test (*p<0,05; **p<0,01; ns, not significant). **H**. Cortical neurons were grown in regular microfluidic chambers for 6 days. Then, TrkB-fc (100 ng/mL) was added to the CB compartment for 60 minutes of treatment. After, the AC was not treated (control) or treated with BDNF (BDNF), or BDNF in the presence of CHX (BDNF + CHX), in the presence of puromycin. **I.** Subsequently, the neurons were fixed, and immunostaining for puromycin and Rab5 was performed, followed by PLA. De novo-synthesized Rab5 protein was observed in green, and the axons were labeled with f-Ctb in red. Scale bar, 10 µm. **J.** Quantification of PLA fluorescence (green) in distal axons for each treatment. n=9-10 compartmentalized chambers from three independent experiments. Values represent the mean ±SD. Statistical analysis was performed by one-way ANOVA and Tukeýs multiple comparison test (****p<0,0001). **K**. Cortical neurons were cultured in radial compartmentalized cultures for 7-8 days. Then, cell bodies were aspirated with a pipette in PBS at 37°C. After 1 hour at 37°C in neurobasal culture media, the remaining axons were untreated (control) or treated for 3 hours with BDNF (BDNF), or BDNF in the presence of CHX (BDNF + CHX). **L.** Then, axonal lysates were obtained, and the levels of NeuN, Rab5, and βIII-tubulin (βIII) were assessed by western blotting. T, total lysates. The levels of Rab5 and βIII from total lysates were developed in separate membranes from the axonal lysates to avoid overexposure. **M.** Quantification of axonal Rab5 normalized by βIII protein levels. Values represent the mean ±SD, n=4. Statistical analysis was performed by one-way ANOVA and Šidák’s multiple comparison test (*p<0,05; **p<0,01; ***p<0,001).

To directly demonstrate BDNF-induced Rab5 translation, we performed proximity ligation assays (Puro-PLA) using antibodies against puromycin and Rab5^34^ (Figure 5H). BDNF significantly increased the Puro-PLA signal for Rab5, an effect that was prevented by CHX (Figure 5I and 5J), providing direct evidence that BDNF stimulates local Rab5 translation in axons.

Finally, we designed an experiment that conclusively demonstrates that BDNF triggers the upregulation of Rab5 in axons independently of the cell body. Using radial microfluidic chambers, we separated axons from their cell bodies by aspirating the cell body compartment with a pipette, then treated axons with vehicle, BDNF alone, or BDNF in the presence of anisomycin (Figure 5K). We prepared axonal lysates and performed western blotting. As shown in Figure 5L and M, BDNF increased Rab5 levels, and this effect was abolished by anisomycin even in the absence of cell bodies.

Collectively, our data demonstrate that cortical axons locally translate Rab5 mRNA into protein in response to BDNF stimulation, in an mTOR-dependent manner. This conclusion is supported by four converging lines of evidence: the BDNF-induced increase in axonal Rab5 protein requires TrkB and mTOR activity; Rab5 mRNA is present in axons; BDNF stimulation increases the puro-PLA signal for Rab5 in axons; and axons stripped of their cell bodies still respond to BDNF with increased Rab5 levels.

### Local translation of Rab5 in cortical neurons is required for BDNF-induced axonal transport and CREB phosphorylation

Given the pivotal role of Rab5 in sorting endosomes into the retrograde transport pathway ^13,15,42^, we hypothesized that BDNF-induced Rab5 synthesis facilitates axonal trafficking. To test this, we developed an approach to specifically knock down Rab5 in axons using siRNA. We first determined that Rab5 has a half-life exceeding 4 hours in cortical neurons (Figure 6A, B), allowing us to design experiments targeting newly synthesized protein. Since our antibody recognizes Rab5A and Rab5B, we used a published siRNA mix targeting both isoforms^43,44^. RT-qPCR confirmed expression of both homologs in cortical neurons (Figure 6C), and the siRNA mix effectively reduced their mRNA levels after 48 hours (Figure 6D). Importantly, Rab5 protein levels measured by immunofluorescence were reduced by approximately 66% relative to control, confirming both antibody specificity and siRNA efficacy (Figures 6E and F). For compartment-specific knockdown, we treated only the AC with siRNA for 4 hours (1 hour pre-treatment, then 3 hours with or without BDNF), while CB contained TrkB-fc. This protocol did not affect cell-body Rab5 levels (Figure 6G-I) but prevented BDNF-induced Rab5 upregulation in axons, without affecting basal Rab5 levels detectable by immunofluorescence (Figure 6J-L).

**Figure 6.**
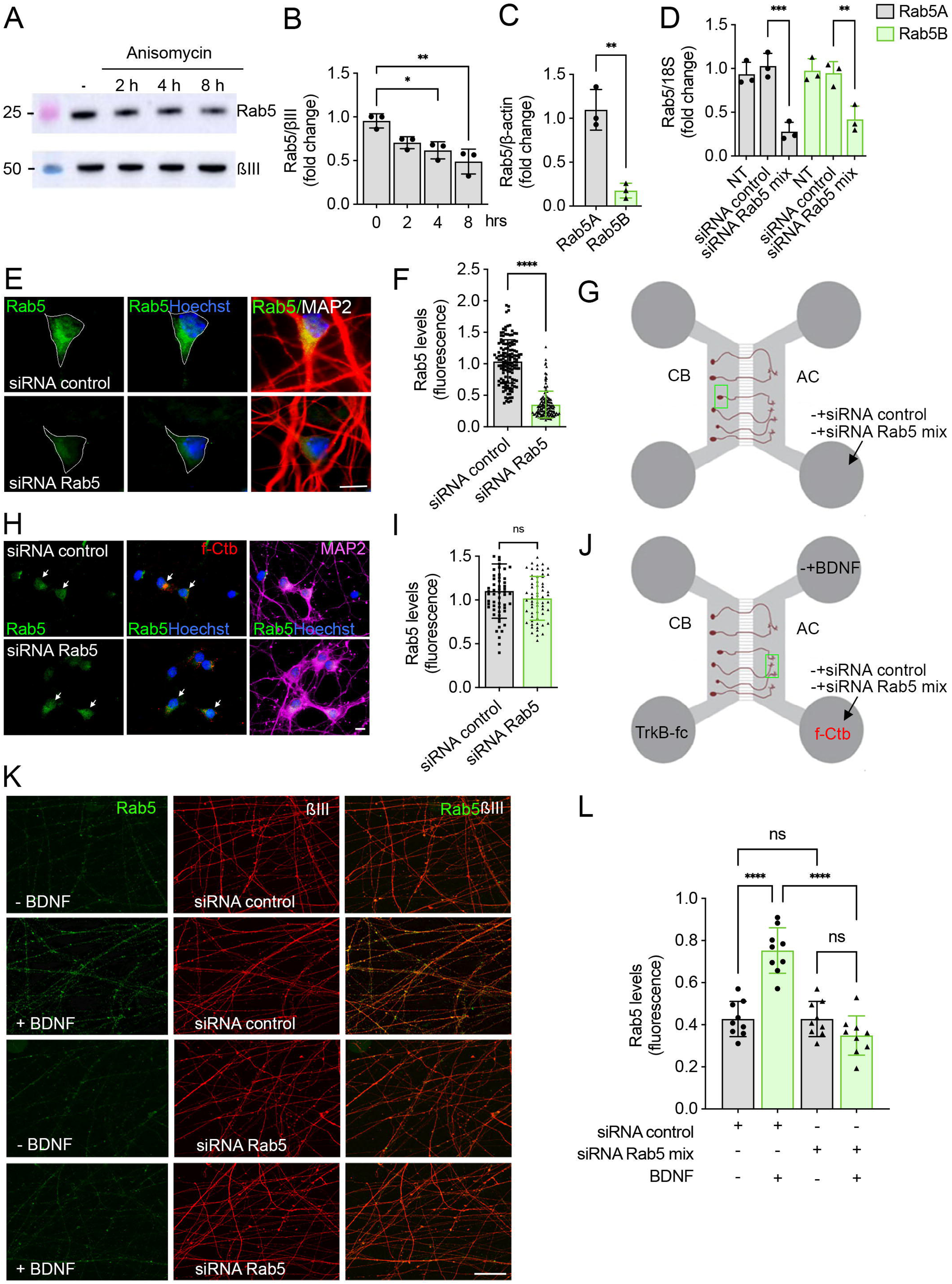
BDNF-induced increase in axonal Rab5 is reduced by axonal treatment of siRNA against Rab5. **A.** Cortical neurons were cultured in p60 plates for 7 days. Then, neurons were either untreated (control) or treated with anisomycin (25 µM) for 2, 4, or 8 hours. The protein levels of Rab5 were studied using western blotting and normalized by βIII protein levels (βIII). **B.** Quantification of Rab5 normalized by βIII protein levels. Values represent the mean ±SD, n=3. Statistical analysis was performed by one-way ANOVA and Bonferronís multiple comparison test (*p<0,05; **p<0,01). **C.** The levels of *Rab5A* and *Rab5B* mRNAs in cortical neurons were studied using RT-qPCR and expressed relative to *Rab5A* levels. *ß-actin* was used as a housekeeping gene. *Rab5A* mRNA levels were about 5-fold higher compared to *Rab5B* mRNA. Values represent the mean ±SD, n=3. Statistical analysis was performed by Unpaired t-test (**p<0,01). **D.** Cortical neurons were cultured in p60 plates for 5 days and then were either untreated (NT), transfected with siRNA control (siRNA control), or with an equimolar mix of siRNA for *Rab5A* and *Rab5B* (siRNA Rab5 mix). After 48 hours, RT-qPCR was performed for *Rab5A* (gray bars) or *Rab5B* (green bars). Results were expressed as fold change over siRNA control treatment. 18S was used as housekeeping. Values represent the mean ±SD, n=3. Statistical analysis was performed by two-way ANOVA and Tukeýs multiple comparison test (**p<0,01; ***p<0,001). **E.** Cortical neurons were cultured in a 12 mm coverslip for 5 days, and then the same treatment was performed as the one described in D. At the end of the treatment, neurons were fixed, and a double immunostaining was performed for Rab5 (green) and MAP2 (red). Nuclei were stained with Hoechst (blue). **F.** The fluorescence levels of Rab5 in neuronal cell bodies were quantified in individual cells and expressed as fold change over control (siRNA control). Values represent mean ± SD of n=135 neurons from three independent experiments. Statistical analysis was performed by Unpaired t-test (****p<0,0001). **G.** Cortical neurons were cultured in regular microfluidic chambers for six days, and the axons (AC) were treated with f-Ctb overnight. At DIV7, siRNA control or an equimolar mix of siRNA for *Rab5A* and *Rab5B* (siRNA Rab5 mix) was added to the AC for 1 hour and then left untreated for 3 hours in complete neurobasal media without antibiotics. **H.** After, the cultures were fixed, and the levels of Rab5 protein (green) and MAP2 (magenta) were visualized by immunofluorescence. Nuclei were stained with Hoechst (blue). **I.** The expression levels of the Rab5 protein were quantified from neuronal cells positive for f-Ctb (red) and MAP2 (magenta). Values represent mean ± SD of n=60 neurons from three independent experiments. Statistical analysis was performed by an unpaired t-test (ns, not significant). **J.** Cortical neurons were cultured and treated as described in G, with the difference that TrkB-fc was added to CB and BDNF was added (or left untreated) to the AC for 3 hours after transfection of axons with siRNAs. **K.** Then, the cultures were fixed, and Rab5 protein (green) and βIII (red) levels were visualized by immunofluorescence in the distal axons. **L.** Quantification of Rab5 immunofluorescence intensity (green) in distal axons standardized by βIII-tubulin (βIII) immunostaining. Values represent the mean ± SD of n=9 chambers from three independent experiments. Statistical analysis was performed using one-way ANOVA and Tukeýs multiple comparison test (****p<0,0001; ns, not significant).

Using this approach (Figure 7A), we found that axon-specific Rab5 knockdown significantly impaired both BDNF-induced f-Ctb retrograde transport (Figures 7B and 7C) and nuclear CREB phosphorylation (Figures 7D and 7E), demonstrating that the newly synthesized Rab5 pool is functionally required for BDNF-enhanced retrograde signaling. Notably, even basal f-Ctb retrograde transport, in the absence of exogenous BDNF, was reduced by Rab5 siRNA, despite no detectable change in steady-state Rab5 immunofluorescence levels (Figures 7B and 7C). This dissociation between total protein abundance and transport capacity indicates that axon terminals are acutely sensitive to the availability of newly synthesized Rab5, and that the pre-existing pool is insufficient to sustain normal trafficking when local synthesis is acutely blocked. Consistent with this, basal nuclear CREB phosphorylation was also reduced (Figures 7D and 7E), suggesting that constitutive retrograde signaling in these cultures similarly depends on ongoing local Rab5 synthesis.

**Figure 7.**
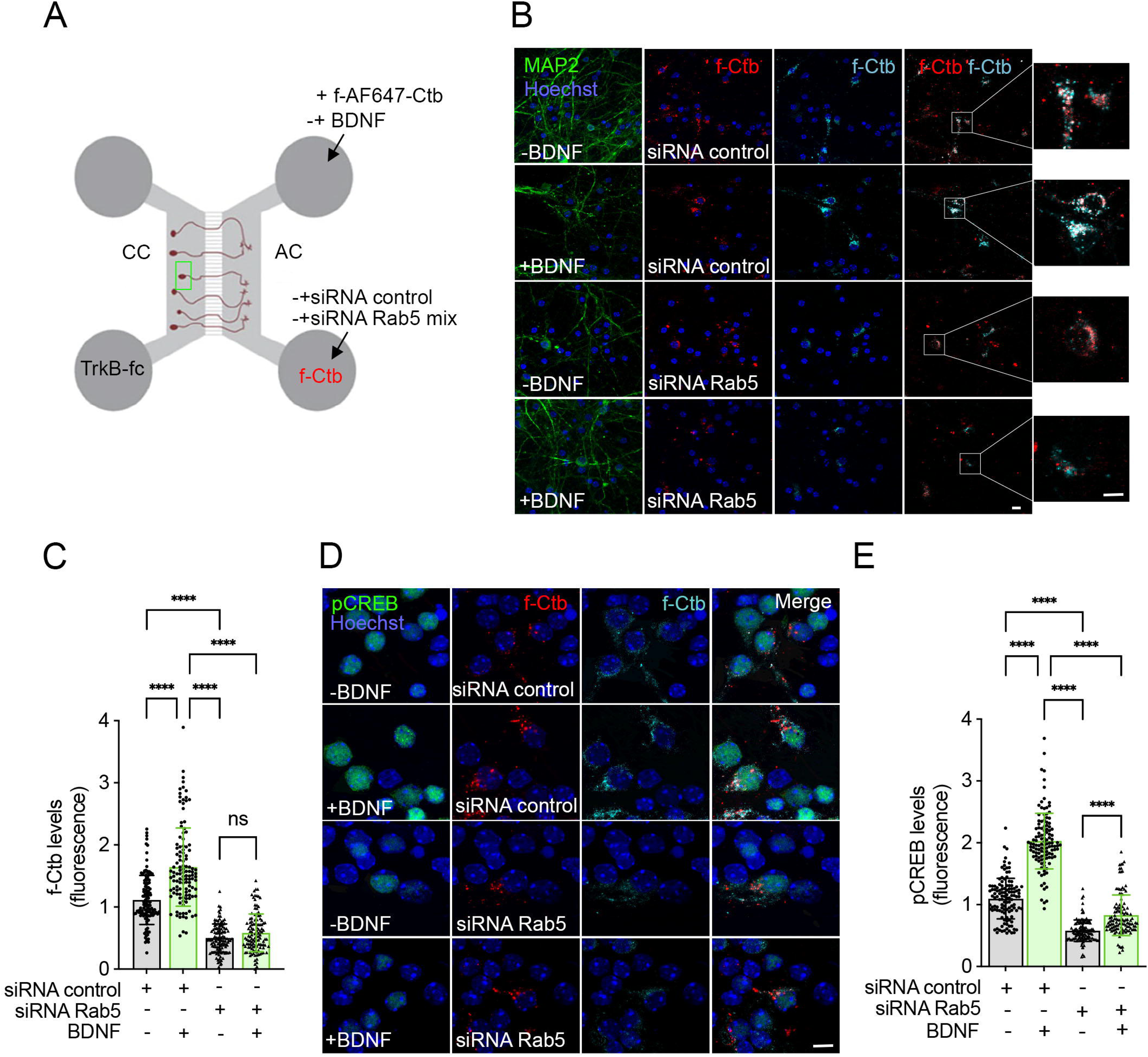
Inhibition of BDNF-induced increase in newly synthesized Rab5, downregulated axonal transport and nuclear responses to axonal BDNF. **A.** Cortical neurons were cultured in regular microfluidic chambers for six days, and the axons (AC) were treated with f-Ctb (f-AF555-Ctb, visualized in red) overnight. At DIV7, siRNA control or an equimolar mix of siRNA for *Rab5A* and *Rab5B* (siRNA Rab5 mix) was added to the AC for 1 hour and then treated for 3 hours with f-Ctb (f-AF647-Ctb, visualized in cyan) in the absence or presence of BDNF. **B.** Then, cultures were fixed and immunostained against MAP2 (green). Nuclei were stained with Hoechst (blue). The axonal transport of f-Ctb was indirectly evaluated in neurons by accumulation of Alexa Fluor 647-Ctb in cell bodies after 3 hours of treatment in axons. Alexa Fluor 647-Ctb was evaluated in neurons positive for MAP2 and Alexa Fluor 555-Ctb. Then, just neurons having axons in the AC were evaluated. Scale bar, 20 μm. Scale bar zoomed up photos, 10 μm. **C.** Quantification of the Alexa Fluor 647-Ctb associated fluorescence intensity in neuronal cell bodies relative to the control. Only cells with an Alexa Fluor 555-Ctb were considered in the quantification. Values represent mean ± SD of n=120 neurons from three independent experiments. Values were normalized to control (siRNA control). Statistical analysis was performed using one-way ANOVA and Tukeýs multiple comparison test (****p < 0.0001, ns, not significant). Scale bar, 10 μm. **D.** Neurons were treated as indicated in A, but after fixation, the cultures were immunostained for pCREB. **E.** Quantification of nuclear pCREB fluorescence intensity. Values represent mean ± SD of 120 cells from three independent experiments. Values were normalized to control (siRNA control). Statistical analysis was performed using one-way ANOVA and Tukeýs multiple comparison test (****p < 0.0001). Scale bar, 10 μm.

Collectively, these results demonstrate that BDNF stimulates mTOR-dependent local translation of Rab5 in axons, and this process is essential for BDNF-induced enhancement of axonal transport and nuclear CREB phosphorylation in cultures of compartmentalized cortical neurons.

## DISCUSSION

Here we demonstrate that axonal BDNF activates mTOR to drive local synthesis of Rab5 at axon terminals, and that this on-demand translational response is required for axonal transport of signaling endosomes and for nuclear CREB activation. These findings reveal a translational layer of axonal BDNF signaling that operates in parallel with, and is mechanistically distinct from, the post-translational mechanisms previously described in this context. PLCγ-Ca² signaling governs internalization of the BDNF/TrkB complex into axonal endosomes^30^, and ERK1/2 modulates dynein subunits to enhance retrograde transport^45^ (Figure 8); however, these pathways act by modifying or recruiting proteins already present at the time of stimulation. mTOR-dependent translation operates differently: rather than remodeling the existing axonal proteome, it expands it by synthesizing new proteins on demand. This distinction implies that the capacity for retrograde signaling is not fixed at the moment of stimulation but can be actively upregulated through local protein translation, a form of plasticity not previously described for axonal BDNF signaling (Figure 8).

**Figure 8.**
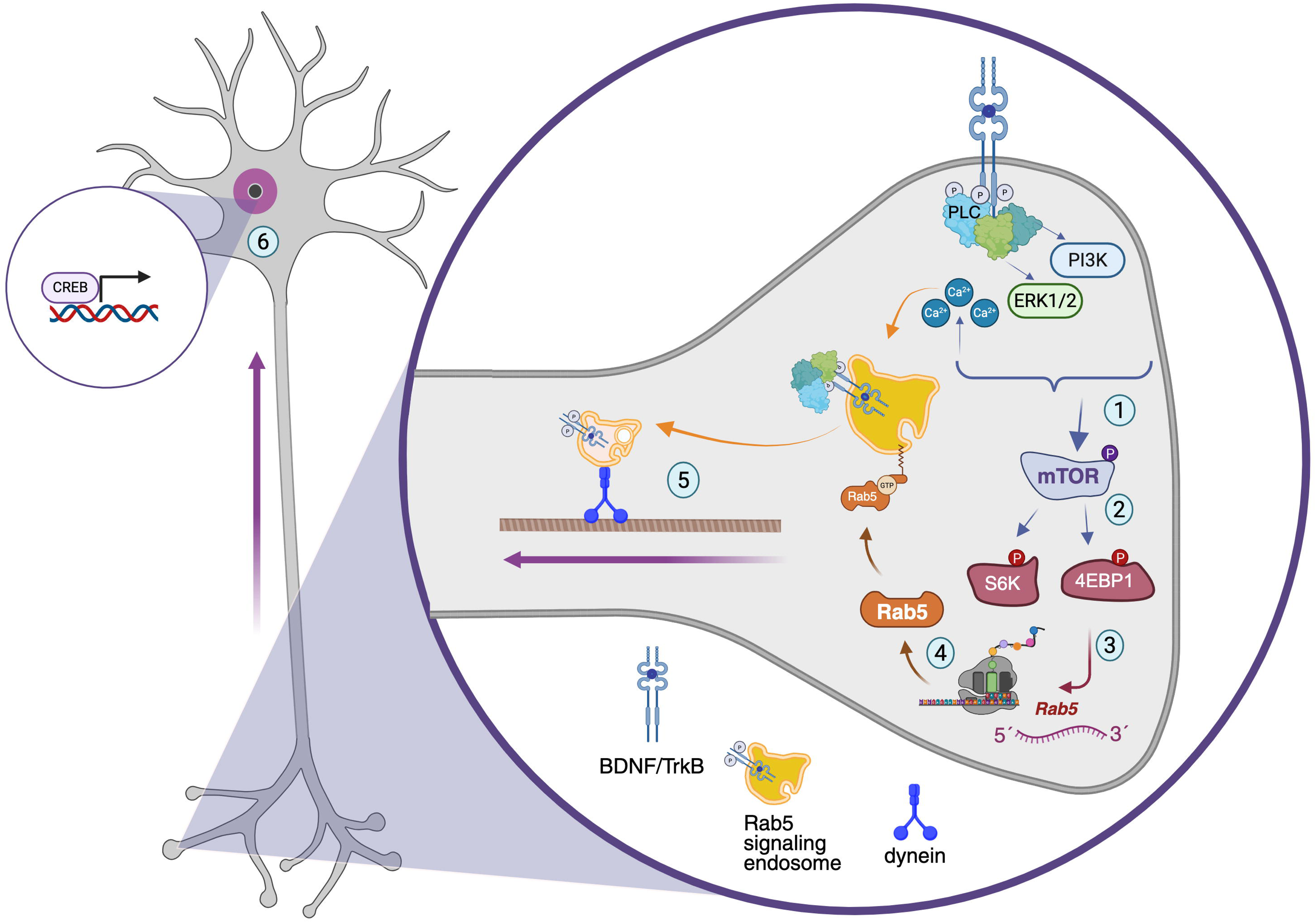
Model for BDNF-induced axonal mTOR-dependent local translation of Rab5 and its role in axon to nucleus communication in cortical neurons. BDNF binding to TrkB at axon terminals triggers distinct signaling pathways that lead to mTOR activation **(1)**. PLCγ (PLC) activation increases TrkB internalization in axons^30^, and ERK1/2 modulates dynein activity^45^. While PI3K-AKT signaling has been shown to result in mTOR activation in postsynaptic boutons^2^, PI3K activity does not play a role in signaling endosomal transport^6^, suggesting that other signaling pathways lead to mTOR activation in axons. Activated mTOR phosphorylates downstream effectors S6K and 4EBP1 **(2)**, which promote the local translation of axonally localized mRNAs, including *Rab5* (3). Newly synthesized Rab5 protein **(4)** is incorporated into the endosomal sorting machinery, where it facilitates the formation of Rab5-positive signaling endosomes containing BDNF/TrkB complexes. These signaling endosomes are coupled to the dynein motor and undergo retrograde transport along axonal microtubules toward the cell body **(5)**. Upon arrival at the soma, retrogradely transported signaling endosomes activate nuclear CREB phosphorylation and CREB-dependent transcription of plasticity-related genes **(6)**. This model reveals that on-demand, mTOR-dependent local synthesis of the trafficking regulator Rab5 in axons is a prerequisite for increasing axonal transport, including the retrograde transport of signaling endosomes. This allows axonal neurotrophic signals to propagate to the nucleus. Created with BioRender.

Local protein synthesis enables neurons to regulate their proteome with spatial and temporal precision in distal compartments. In the peripheral nervous system, axonal translation is well established and plays key roles in injury responses and regeneration^22,46^. In CNS axons, the picture has been slower to emerge, partly because early studies found limited ribosomal content in mature central axons, suggesting that they function as single ribosomes, or monosome^47^. This, together with more recent evidence, has overturned this view: axonal mRNA populations have been characterized in compartmentalized cortical neuron cultures^48^, β-actin translation is required for arborization of retinal ganglion cells^49^, and active translation in the visual circuit has been demonstrated in vivo during both development and adulthood^50^. Beyond growth cones, local protein synthesis in presynaptic compartments is now recognized as a widespread feature of the adult brain^51^, with functional roles in presynaptic terminal formation^52^ and NMDA receptor-dependent sustained firing^53^.

Notably, mTOR activity has been linked to axonal regeneration and integrity^54–56^, and growth cone proteomes from callosal axons are enriched in mTOR kinase and mTOR-dependent transcripts^57^, suggesting that the mTOR-translation axis is an intrinsic feature of axonal biology rather than a response exclusive to injury or development. Our findings extend this framework to cortical neuron axons, showing that BDNF-activated mTOR drives on-demand Rab5 synthesis and couples local proteostasis to retrograde signaling and nuclear CREB activation.

The observation that BDNF enhances retrograde transport of both signaling endosomes and f-Ctb^6,30^ suggests that mTOR-dependent translation broadly activates axonal endosomal pathways rather than selectively promoting a single cargo, consistent with earlier findings that elevated axonal calcium enhances transport of both TrkB and f-Ctb^37^. This points to a model in which newly synthesized proteins, including but not limited to Rab5, collectively increase the capacity of the axonal endosomal system to sort, form, and sustain retrogradely transported carriers. In this framework, Rab5 serves as a rate-limiting node: its local synthesis is necessary but likely not sufficient, and identifying the full complement of mTOR-dependent axonal proteins that cooperate with Rab5 to support retrograde transport will be an important goal for future studies. Axon-specific ribosome profiling approaches, using RiboTag or proximity-labeling strategies, are well-suited for this purpose.

The reduction in basal retrograde transport caused by axon-specific Rab5 siRNA, despite no detectable change in steady-state Rab5 immunofluorescence, is mechanistically informative. Given that Rab5 has a half-life exceeding four hours, this effect cannot be attributed to bulk protein degradation during the siRNA treatment window, suggesting instead that newly synthesized Rab5 plays a functionally distinct role that the pre-existing pool cannot substitute. One possibility is that newly synthesized Rab5 is preferentially targeted to nascent endosomes or to specific membrane subdomains required for the initial steps of endosome biogenesis, a role that aged or membrane-resident Rab5 may be unable to fulfill. The correlated reduction in basal nuclear CREB phosphorylation further indicates that constitutive retrograde signaling depends on ongoing Rab5 synthesis. Together, these findings suggest that local translation is not simply an amplification mechanism engaged under strong stimulation but a constitutive feature of axonal homeostasis, with significant implications for how we understand the maintenance of long-distance neuronal communication under baseline conditions.

How the translational response is spatially restricted to axons remains an important open question. Since BDNF activates mTOR in both axons and dendrites, spatial specificity cannot be determined solely at the level of kinase activation. A key upstream determinant is the prior localization of specific mRNAs: Rab5 mRNA must be present in axons before it can be translated there, a prerequisite we confirm using RT-qPCR of compartment-specific RNA fractions (Figure 5A–C). mRNA localization is mediated by cis-regulatory elements in the 3’UTR, recognized by RNA-binding proteins (RBPs) that form translationally repressed ribonucleoprotein complexes, which are mobilized in response to local signals, including neurotrophin stimulation^58–60^. While the specific RBPs and 3’UTR elements governing axonal Rab5 mRNA localization are not yet known, the 3’UTRs of other locally translated GTPases, such as RhoA and β-actin, contain well-characterized zipcode elements^27,40,41^, suggesting that analogous mechanisms may operate for Rab5. Identifying these elements would enable selective manipulation of axonal Rab5 translation without affecting somatic expression, a tool of both mechanistic and therapeutic value.

In both mice and humans, Rab5 exists as three homologs, Rab5A, Rab5B, and Rab5C, encoded by separate genes^61^. While these isoforms share core roles in early endocytic transport, they also exhibit unique functions in non-neuronal contexts^62–65^, and their combined depletion leads to the loss of early and late endosomes and cell death^43,66^. Whether Rab5 isoforms perform redundant or distinct functions in neurons remains unknown. Given this, our study employed a combined Rab5A/Rab5B knockdown strategy to assess their collective contribution to axonal mTOR-dependent local translation, without presupposing isoform-specific roles that the existing literature does not yet support.

Our findings suggest a previously unrecognized link between axonal proteostasis and neurodegenerative disease. In ALS, axonal TDP-43 aggregation suppresses local synthesis of nuclear-encoded mitochondrial proteins^39^, and C9ORF72 haploinsufficiency disrupts endocytic pathways in motor neurons^67^. In Alzheimer’s disease, enlargement of Rab5-positive early endosomes is among the earliest pathological changes, preceding amyloid plaque deposition^68,69^. If constitutive local Rab5 synthesis is required for basal retrograde transport, as our data suggest, then impairment of axonal translation or mTOR signaling could directly compromise BDNF retrograde signaling and neuronal survival. This would position axonal proteostasis as an upstream contributor to transport deficits in neurodegeneration and may help explain why axonal transport failure and translation disruption are early, convergent features of multiple neurodegenerative conditions. These connections await validation in disease models, but provide a conceptual framework linking axonal proteostasis to BDNF-dependent neuronal maintenance.

Several important questions remain. The upstream mechanism linking axonal TrkB activation to mTORC1 is not yet defined. In dendrites and cell bodies, the canonical pathway involves PI3K-AKT downstream of TrkB, leading to TSC1/2 inhibition and mTORC1 activation^18^. However, in our previous work^6^, inhibition of PI3K in axons did not preclude signaling endosome transport, suggesting that the classic PI3K-mTOR pathway is not involved. Whether alternative routes, such as ERK1/2 or PKC^29,70^, lead to axonal mTOR activation downstream of BDNF remains to be determined. The spatial constraints of the axon may impose different requirements on signal amplification and kinase compartmentalization. Beyond Rab5, the full mTOR-dependent axonal translatome has yet to be mapped; axon-specific ribosome profiling will be essential for identifying additional locally translated effectors. Finally, whether BDNF-induced axonal Rab5 translation operates in vivo during learning and memory, where BDNF-dependent synaptic plasticity is required, is a critical question that will require combining axon-specific translation reporters with behavioral paradigms and circuit-level analysis.

## METHODS

### Materials

List of Materials in Supplementary Material.

### Isolation and Culture of Embryonic Cortical Neurons

Embryonic cortical neurons were isolated from C57Bl/6 J or TrkB^F616A^ knock-in mice ^38^ on embryonic days 16–17 and obtained from the animal facilities of our institution. Pregnant animals were euthanized under deep anesthesia following bioethical protocols approved by the Bioethics Committee of the Universidad Andrés Bello (009/2022).

### Preparation of neuronal cultures

Cortical tissues from mice were dissected and dissociated into single cells in Hank’s balanced salt solution (HBSS). After dissociation, the neurons were resuspended in Modified Dulbecco’s Modified Eagle Medium (DMEM) supplemented with 10% horse serum (HS) (DMEM/HS) and seeded in microfluidic chambers at a density of 50,000 cells/microfluidic chamber. Three hours later, the culture medium was replaced with complete neurobasal supplemented with 2% B27, 1x GlutaMAX, and 1x penicillin/streptomycin, as well as 0.25 μM cytosine arabinoside (Ara-C)^6^.

### Compartmentalized cultures

The molds used to prepare standard two-chamber microfluidic cultures, as previously published ^6,30^, and radial compartmentalized chambers, as published in Altman et al. 2021^39^ were fabricated at the microfluidic core facility of Tel Aviv University^71^. Microfluidic chambers were prepared using a SyldgardTM 184 silicone elastomer base, following the manufacturer’s instructions. Two days before primary culture, glass coverslips (25 mm for the standard microfluidic chamber or 12 mm for the radial microfluidic chamber) were incubated overnight with poly-D-lysine (0.1 mg/mL). The next day, the poly-D-lysine was washed with MilliQ-ultrapure water, and microfluidic chambers were placed on the dry coverslips. Then, laminin (2 μg/mL in MilliQ ultrapure water) was added to the chamber overnight. On the same day of the primary culture, the laminin solution was replaced with DMEM/HS (DMEM, supplemented with 10% Horse Serum, 1x GlutaMAX, and penicillin/streptomycin).

Standard two-chamber compartmentalized cultures were seeded at a density of 50,000 cells in the cell body compartment in 80 μL of DMEM/HS, and 160 μL of the medium was added to the axonal compartment. This procedure prevented cells from migrating to the axonal compartment. DMEM/HS is replaced three hours later by 160 µL of complete Neurobasal medium in the cell body compartment and 80 µL of complete Neurobasal medium in the axonal compartment. The volume difference between the two compartments must be at least 80–100 µL to ensure microfluidic isolation. This differential volume must be conserved throughout all cultures if fluidic isolation is required and to prevent evaporation. Every 48 hours, the culture medium is replaced with fresh, complete neurobasal medium, preserving the volume differential between the two compartments. Experiments are performed on day in vitro seven (DIV7).

For compartmentalized cultures in radial microfluidic chambers, 12-mm coverslips are placed in 24-well plates, treated with poly-D-lysine (0.1 mg/mL), and coated with laminin (2 μg/mL) as described for standard microfluidic chambers. Neurons were seeded at a density of 100,000 cells diluted in 60 µL of DMEM/HS and added to the central cell body compartment. On the other hand, 400 μL of DMEM/HS was added to the axonal (outside) compartment. After 3 hours, the DMEM/HS was replaced with complete neurobasal medium by adding 400 μL of medium to the axonal compartment and 100 μL to the cell body compartment. The media was partially refreshed every 48 hrs. Compartmentalized cultures were used at DIV7 for RNA extraction to detect mRNA using RT-qPCR and for biochemical analyses of protein after BDNF treatment.

### Axonal treatment with BDNF

Each two-chamber compartmentalized culture was evaluated for morphological and fluidic compartmentalization by using an axonal treatment with recombinant cholera toxin subunit B conjugated to Alexa Fluor 555 (f-Ctb) at a final concentration of 1 μg/mL. f-Ctb was added to the axonal compartment at DIV6 and incubated overnight. Experiments were conducted at DIV7. The cell media were removed from each compartment, and 160 µL of warm neurobasal medium containing reduced 0,5% B27 (reduced neurobasal medium), supplemented with TrkB-fc (100 ng/mL), was added to the cell body compartment. To the axonal compartment, 80 μL of warm reduced neurobasal medium was added and incubated for 1 hour at 37 °C. Then, the media were replaced, specifically in the axonal compartment, with 80 μL of reduced neurobasal medium alone (control) or supplemented with BDNF at 50 ng/mL, in the presence or absence of inhibitors for 180-minute treatments. Afterward, the drugs were removed and washed twice with 120 μL of PBS supplemented with protease and phosphatase inhibitors. A similar protocol was followed for radial chambers; the inner compartment (cell bodies) contained neurobasal medium with TrkB-fc, and the outer compartment contained reduced neurobasal medium alone (control) or BDNF-supplemented neurobasal medium.

### Axonal immunostaining

To fix neurons in a two-chamber compartmentalized culture, the chamber was carefully removed, and 200 µL of a 4% paraformaldehyde (PFA) and 4% sucrose solution diluted in PBS (PFA solution) was added to the coverslip, which was then incubated at room temperature (RT) for 18 min. Then, the PFA solution was removed, and three washes (5 minutes each) with 150 µL RT PBS were performed. Neurons were permeabilized with 150 μL of blocking solution (3% bovine serum albumin, IgG and protease-free, BSA, 5%, fish gelatin, 0,2% saponin in PBS) for 60 min at RT. Appropriately diluted primary antibodies were incubated in 100 μL of antibody solution (BSA 3%, fish gelatin 5%, saponin 0,02% in PBS) overnight at 4°C. The dilution for the first antibodies was as follows: rabbit anti-Rab5 (1:250), rabbit anti-phospho S6 (1:300), rabbit anti-phospho 4EBP1 (1:250), rabbit anti-phospho CREB (1:400), mouse anti-βIII-tubulin (1:750), and mouse anti-MAP2 (1:750). The next day, the primary antibody was removed, and three PBS washes were performed, each for 5 minutes. The PBS buffer was removed, and neurons were incubated with 100 µL of secondary antibody solution for 90 min at RT. Donkey anti-rabbit Alexa 488 (1:500) and anti-mouse Alexa 647 (1:500) were used as secondary antibodies in the experiments. The secondary antibody was removed, and coverslips were washed with cold PBS for 5 minutes. Then, samples were incubated with 100 µL of Hoechst solution (1:5000) for 10 min. The solution was removed, and cells were rewashed with cold PBS, then with cold distilled water, to remove salts. Finally, the coverslip was mounted in Mowiol.

### Axon-specific siRNA transfection in compartmentalized microfluidic cultures

Sequences for the siRNA control and for siRNA targeting Rab5A and Rab5B mRNA were obtained from ^43,44^ and synthesized by Integrated DNA Technologies (IDT). Compartmentalization was verified by adding f-Ctb (1 μg/mL) to the axonal compartment overnight at DIV6. At DIV7, conditioned medium was removed from the axonal compartment and replaced with antibiotic-free neurobasal medium for 1 hour. Simultaneously, TrkB-fc (100 ng/mL) was added to the cell body compartment and maintained throughout the entire treatment period to sequester endogenous BDNF. Transfection complexes were prepared by separately diluting 0.2 μL of Lipofectamine RNAiMAX and 1.5 pmol of siRNA (siRNA control or siRNA Rab5A+B mix) in 10 μL of Opti-MEM, then mixing and incubating at room temperature for 10 minutes. Twenty microliters of the transfection mixture were added to each axonal compartment and incubated for 1 hour at 37°C. Following transfection, BDNF (50 ng/mL) was added to the axonal compartment in the presence of siRNA, and cultures were incubated for 3 hours. Microfluidic chambers were then carefully removed, and cells were fixed with 200 μL of 4% PFA and 4% sucrose in PBS for 18 minutes at room temperature and processed for immunofluorescence as described above.

### Axonal phospho-S6 (pS6), phospho-4EBP1 (p4EBP1), and anti-βIII-tubulin protein quantification from cortical neurons by confocal microscopy

Three independent experiments were conducted (each corresponding to a different primary culture), and three compartmentalized chambers were obtained for each experimental condition. Then, three fields per region of the axonal compartment (microgrooves, proximal, and distal) were captured at 63X magnification using a Leica SP8 confocal microscope in each compartmentalized chamber. To obtain the maximum-fluorescence-intensity projection, 5- to 8-stack images with 1-μm spacing were acquired. The fluorescence intensity per field was calculated by applying a threshold in ImageJ (FIJI version 1.5) to remove background from the microphotograph, and the resulting image was then used to quantify the fluorescence associated with pS6 and p4EBP1. This value was standardized to βIII-tubulin levels by dividing it by the fluorescent intensity obtained from the βIII-tubulin channel. The average of the three fields for each region (microgrooves, proximal, and distal) was calculated to yield the fluorescent intensity per region per chamber. The quantification was performed using the FIJI ImageJ version 1.5 software, and the protein of interest was plotted and standardized by βIII-tubulin levels.

### Quantification of BDNF-induced retrograde transport of f-Ctb

Neurons with axons in the axonal compartment (AC) were labeled by treating the AC with 1ug/ml f-Ctb in the presence or absence of BDNF (50 ng/ml) for 180 minutes. The effect of local translation for BDNF-induced retrograde transport of f-Ctb was assessed by treating the axons with anisomycin (25 μM). This treatment was initiated in the AC 1 hour before adding BDNF, simultaneously with depletion of endogenous BDNF in the CB using TrkB-fc, and continued for the next 180 minutes of BDNF treatment. Subsequently, the f-Ctb label is washed, and the cultures are fixed as indicated above. Mouse anti-MAP2 (1:750) was used as a neuronal marker. The cells were observed under epifluorescence microscopy.

In the case of the quantification of axonal transport of axons treated with siRNAs an additional consideration was taken. At DIV6, the AC of microfluidic cultures were treated with cholera toxin subunit B conjugated to Alexa Fluor 555 (f-AF555-Ctb) at a final concentration of 1 μg/mL and left ON. Then, at DIV7, the BDNF treatment was performed in the presence of cholera toxin subunit B conjugated to Alexa Fluor 647 (f-AF647-Ctb) at a final concentration of 1 μg/mL. The cells were observed using a Leica SP8 confocal microscope.

The intensities of the f-AF555-Ctb (in experiments with anisomycin) or f-AF647-Ctb (in axonal siRNA treatments) fluorescence signals were initially quantified in ImageJ by subtracting the background autofluorescence of the label. This background was determined by measuring the intensity of images captured with a 555- or 647-nm filter under experimental conditions in the absence of f-Ctb. After background subtraction, regions of interest (ROIs) were generated for cells labeled with f-Ctb. Only cells containing more than three Ctb vesicles were included in the analysis. The selection tool was then used to delineate Ctb fluorescence areas, and fluorescence intensities were measured for 45–50 cells per condition. The average Ctb intensity of the control condition was calculated, and each fluorescence intensity value from the experimental condition (+BDNF) was normalized by dividing it by the average intensity of the control condition.

### Determination of phospho-CREB (pCREB) levels in the nucleus of compartmentalized cortical neurons

Neurons with axons in the axonal compartment (AC) were labeled by treating axonal compartments ON with f-Ctb (Figure 1A and B). Then, the AC was treated with BDNF (50 ng/mL) in the presence or absence of cycloheximide (25 µM) for 180 min in the axonal compartment. After fixing the cultures, they were immunolabeled for pCREB (Ser 133). The fluorescence intensity of 15 cells per chamber labeled with f-Ctb was measured in three independent experiments, with two chambers per experiment.

In the case of the quantification of pCREB levels in cultures where axons were treated with siRNAs, the same consideration was taken as indicated above for axonal transport of f-Ctb. At DIV6, the AC of microfluidic cultures was treated with f-AF555-Ctb ON. Then, at DIV7, the BDNF treatment was performed in the presence of f-AF647-Ctb. After fixing the cultures, they were immunolabeled for pCREB (Ser 133). The fluorescence intensity of 20 cells per chamber labeled with f-Ctb was measured in three independent experiments, with two chambers per experiment. Nuclei were stained with Hoechst and observed using a Leica SP8 confocal microscope.

### O-propargyl-puromycin (OPP) protein synthesis assay

Standard compartmentalized cultures were used at DIV 7. The complete neurobasal medium was removed from the cell body compartment and replaced with reduced neurobasal medium supplemented with TrkB-fc (100 ng/ml) for 60 min. In the last 30 minutes of this treatment, the complete neurobasal medium was replaced from the axonal compartment by reduced B27 neurobasal medium in the presence or absence of 25 μM cycloheximide (CHX) or 0.25 μM Torin1 for the remaining 30 min. Subsequently, the medium was removed from the axonal compartment and replaced with reduced B27 neurobasal medium supplemented with 20 μM OPP, with or without BDNF (50 ng/mL), and with or without CHX or Torin1 inhibitors, and the mixture was incubated for 60 min at 37°C. The medium was removed, and neurons were washed with RT PBS supplemented with protease inhibitors (100 uL/cover). Then, cells were fixed by adding RT PFA solution for 18 min. After removing PFA, the cells were washed twice with ice-cold PBS supplemented with protease inhibitors. Covers were incubated for 60 min in blocking solution, washed with ice-cold PBS, and then incubated with 50 μL of OPP detection mix based on click-iT chemistry according to the manufacturer’s instructions. The cover was washed with 150 uL of cold PBS. The axonal tubulin was labeled with anti-ßIII tubulin (1:750) in antibody solution overnight at 4°C, and the procedure was then continued as described above for axonal immunostaining.

### Determination of axonal mRNA

Cortical neurons were cultured in radial microfluidic chambers. For this procedure, cells received no treatment. RNA from axons and cell bodies was extracted from the outer and inner compartments of the radial microfluidic chambers, respectively, at DIV7, using the RNeasy kit following the instructions of the manufactures. Ten chambers were required for each condition. Reverse transcription and qPCR were subsequently performed using the SuperScript™ First-Strand Synthesis System for RT-PCR with Oligo dT primers, according to the protocol of the manufacturer. Real-Time PCR was conducted to analyze the mRNA levels of polymerase B (PolB) (Fw: ACTTCACGTCAGAATCCAGC; Rv: CCTTTCCATCCTTCTCGCTG), Beta-actin (Fw: ACCTTCTACAATGAGCTGCG; Rv: CTGGATGGCTACGTACATGG), and Rab5 (Fw: CTATGCAGATGACAACAGCTT; Rv: TCAGTTACTACAACACTGGCT). Additionally, the ribosomal 18S gene (Fw: AGTCCCTGCCCTTTGTACACA; Rv: CGATCCGAGGGCCTCACTA) was used as a housekeeping gene.

For RNA/protein biochemical analyses, it must be noted that a large volume difference between the inner (CB) and outer (AC) compartments must be maintained during lysis. Lysis of the outer compartment (axonal) is typically started first. This is achieved by overfilling the inner well with medium/PBS and completely draining the AC just before the lysis reagent is applied. After the lysate has been removed from the outer compartment, the radial ring can be removed, and the lysis reagent can be applied to lyse the CB.

### Biochemical analyses of cell body and axonal proteins after BDNF treatment

To study axonal Rab5 protein levels and other cellular markers by western blot, cortical neurons grown in radial microfluidic chambers were used. The protocol for axonal stimulation with BDNF and drug treatments was the same as that described for standard microfluidic cultures. The cell body compartment (CB) was maintained with TrkB-fc, and the axonal compartment (AC) was treated with or without BDNF, CHX, or anisomycin. Ten radial chambers were pooled per condition. To prepare CB and AC lysates for western blot analysis, radial chambers grown in 24-well plates at DIV7 were washed with cold PBS and placed on ice. Axons were lysed first by adding 100 µL of ice-cold RIPA buffer (10mM Tris-HCl pH 8.0, 150 mM sodium chloride, 1mM EDTA, 0,5% sodium deoxycholate, 0,5% NP40, 0,1% sodium dodecyl sulfate, and a mix of protease and phosphatase inhibitors) to the outer compartment. The radial ring was then removed, and cell bodies were lysed using the same buffer. Lysates from 10 chambers were pooled per condition to achieve sufficient protein concentration for detection.

To study axonal responses without the participation of cell bodies, we also used radial chambers, and axons were separated from their cell bodies by aspirating the cell body compartment with a pipette, then treated with vehicle, BDNF alone, or BDNF in the presence of anisomycin. Lysates from 12 chambers were pooled per condition to achieve sufficient protein concentration for detection. Protein concentration from axonal samples was quantified, and 40 µg of protein per line was loaded onto 12% SDS-PAGE gels. A non-compartmentalized total neuronal lysate was included as a control. Proteins were separated by SDS-PAGE at 90 V for 3 hours and subsequently transferred onto a nitrocellulose membrane overnight at 4°C at a constant current of 100 mA. Membranes were then blocked in 5% non-fat milk in TBS-T (20mM Tris, 150mM sodium chloride, 0,1% tween 20) for 1 hour at room temperature and incubated overnight at 4°C with primary antibodies against Rab5 (1:1000), βIII-tubulin (1:4000), NeuN (1:1000), Lamin (1:300), and Tau (1:1000). The following day, membranes were washed with TBS-T and incubated with the appropriate HRP-conjugated secondary antibodies for 2 hours at room temperature. After washing, membranes were developed using an enhanced chemiluminescence detection reagent, and signals were captured using the LAS-500 imaging system with exposure times of up to 2 minutes. Band intensities were quantified from the acquired images using ImageJ software.

### Determination of Puro-PLA for Rab5

Mouse cortical neuron in regular compartmentalized cultures was treated with f-Ctb and BDNF as previously described. Puromycin (20 µM), with or without CHX (25 µM), was added to axons for 60 minutes before BDNF treatment. Following this, axons were exposed to BDNF (50 ng/mL) or its absence for 180 minutes, with the previously added drugs maintained throughout the treatment. After the treatments, all reagents were removed, and the cultures were washed with cold PBS. The cells were then fixed and permeabilized as described in the Immunostaining section.

Proximity Ligation Assay (PLA) was performed using the NaveniFlex™ Cell MR kit the Navinci, with rabbit anti-Rab5 and mouse anti-puromycin antibodies, incubated overnight at 4°C. Coverslips were subsequently mounted on slides with 20 µL of Mowiol. De novo-synthesized Rab5 was visualized as dots after subtracting the background signal from a negative control lacking the primary antibodies. The number of dots was quantified using a Leica SP8 confocal microscope at 63× magnification, acquiring maximum projections from 5–8 stacks with 1 µm separation.

Dot quantification per chamber was performed using FIJI ImageJ software with the “Analyze Particles” plugin, considering dots larger than 0.3 µm in diameter.

### Determination of the half-life of Rab5 protein and mRNA levels for Rab5a and Rab5b

Seven DIV Cortical neurons were seeded at a density of one million cells per dish on p60 plates coated overnight with poly-L-lysine and maintained in 1.5 mL of complete neurobasal medium. To determine the half-life of Rab5 protein, neurons were processed and lysed with RIPA buffer (control) or treated with anisomycin for 2, 4, or 8 hours to inhibit protein translation and then lysed with RIPA buffer. Then, Rab5 protein was detected by SDS-PAGE followed by western blotting as described for axonal Rab5, except that 10 ug was loaded per lane.

To detect mRNA of *Rab5A*, *Rab5B*, and 18S in non-treated neurons or treated with siRNAs (siRNA control, siRNA Rab5A, siRNA Rab5B, or a mix) as described below, neurons were lysed with TRIzol^®^ reagent to extract RNA. Reverse transcription and qPCR were subsequently performed using the iScript™ cDNA Synthesis Kit and Brilliant II SYBR green qPCR master mix Kit, according to the protocol of the manufacturer. Real-Time PCR was conducted to analyze the mRNA levels of Rab5A (Fw: TGGTTCTTCGCTTTGTGAAAGG; Rv: GCTATGATACCGTTCTTGACCAG), Rab5B (Fw: CAGGCTGCAATCGTGGTCTAT; Rv: ATTCGTGGGTTCGCTCTTTGG). Additionally, the β*-actin* mRNA (Fw: ACCTTCTACAATGAGCTGCG; Rv: CTGGATGGCTACGTACATGG) and the ribosomal 18S gene (18S) (Fw: AGTCCCTGCCCTTTGTACACA; Rv: CGATCCGAGGGCCTCACTA) were used as housekeeping genes.

### siRNA knockdown efficiency validation in dissociated cortical neuron cultures

Cortical neurons in p60 plates or 24-well plates were used to validate siRNA knockdown efficiency. At DIV7, conditioned medium was collected and replaced with antibiotic-free neurobasal medium for 1 hour. Transfection complexes were prepared using Lipofectamine RNAiMAX and Opti-MEM. For p60 dishes, 10 μL of Lipofectamine RNAiMAX and 75 pmol of siRNA (siRNA control or siRNA Rab5A+B mix, consisting of equimolar amounts of siRNA Rab5A and siRNA Rab5B at 10 μM stock) were each diluted in 500 μL of Opti-MEM, mixed, and incubated for 10 minutes at room temperature. 1 mL of the transfection mixture was added per dish, and the dishes were incubated for 2–8 hours at 37°C. For 24-well plates, the same protocol was applied with volumes scaled proportionally (1 μL Lipofectamine RNAiMAX, 7.5 pmoles siRNA, 50 μL Opti-MEM per reagent, 100 μL mixture per well, 4-hour incubation). Following transfection in both formats, conditioned medium was returned, and cells were maintained for 48 hours before RNA extraction (p60 dishes) or immunofluorescence processing (24-well plates).

### Statistical Analysis

The results are expressed as the averages ± standard deviation (SD) or ± standard error of the mean (SEM), depending on the variability of the data. Student’s *t-*test or one-way ANOVA followed by the appropriate multiple comparisons test was performed, depending on the number of groups used in each experiment. The details of the specific test used, the level of significance, and the number of replicates are indicated in each figure legend. Statistical analyses were performed using GraphPad Prism 10 (Scientific Software).

## Acknowledgement

The authors gratefully acknowledge financial support from ANID (Agencia Nacional de Investigacion y Desarrollo), FONDECYT grant no. 1221203 to FCB, FONDECYT grant no. 1220480 to GA. DI-04-24/NUC from Universidad Andres Bello to G.A. and FCB. FONDECYT postdoctoral grant no. 3200800 and a postdoctoral grant from Secretaria de Educación, Ciencia, Tecnología e Innovación de la Ciudad de México (SECTEI) sectei/113/2022 to RTF. DI-10-25/ATP grant from Universidad Andres Bello to PAM.

## Notes

### Competing Interest Statement

The authors have declared no competing interest.

### Summary of Updates

The manuscript has been updated to include new data supporting the role of newly synthesized Rab5 in axons as a central protein in BDNF axonal signaling. Supplementary Figure S1 has been removed from the manuscript, as it is redundant. Other figures have been consolidated into two, and new Figures have been added to the manuscript, including one summarizing our data.

